# Native or non-native protein-protein docking models? Molecular dynamics to the rescue

**DOI:** 10.1101/2021.04.02.438171

**Authors:** Zuzana Jandova, Attilio Vittorio Vargiu, Alexandre M. J. J. Bonvin

**Affiliations:** Computational Structural Biology Group, Bijvoet Centre for Biomolecular Research, Faculty of Science - Chemistry, Utrecht University, Padualaan 8, 3584 CH Utrecht, the Netherlands; Physics Department, University of Cagliari, Cittadella Universitaria, S.P. 8 km 0.700, 09042 Monserrato, Italy

**Keywords:** Protein-protein docking, native pose identification, molecular dynamics simulations, machine-learning, HADDOCK, GROMACS

## Abstract

Molecular docking excels at creating a plethora of potential models of protein-protein complexes. To correctly distinguish the favourable, native-like models from the remaining ones remains, however, a challenge. We assessed here if a protocol based on molecular dynamics (MD) simulations would allow to distinguish native from non-native models to complement scoring functions used in docking. To this end, first models for 25 protein-protein complexes were generated using HADDOCK. Next, MD simulations complemented with machine learning were used to discriminate between native and non-native complexes based on a combination of metrics reporting on the stability of the initial models. Native models showed higher stability in almost all measured properties, including the key ones used for scoring in the CAPRI competition, namely the positional root mean square deviations and fraction of native contacts from the initial docked model. A Random Forest classifier was trained, reaching 0.85 accuracy in correctly distinguishing native from non-native complexes. Reasonably modest simulation lengths in the order of 50 to 100 ns are already sufficient to reach this accuracy, which makes this approach applicable in practice.

## INTRODUCTION

Modelling molecular processes in living organisms is a challenging endeavour in every aspect. Ideally, one would have to capture every component of the molecular machinery throughout all states of their biological journey. It is however already computationally highly demanding to process only a fraction of those interacting pathways in full detail; thus, one has to compromise on the level of detail and/or the size of the simulated system. Docking is a molecular modelling approach commonly used to target large-scale interactions of two or more binding partners of any nature. It proficiently explores the possible binding modes throughout the conformational/interaction space, what is referred to as sampling. A number of software packages like HADDOCK,(1,2) LightDock,(3,4) ATTRACT,(5,6) IMP(7,8) or ROSETTA(9,10) allow to efficiently utilise available experimental and/or bioinformatics information to eliminate sampling of irrelevant binding modes and guide protein-protein docking to meaningful outcomes. The next step is to detect the most favourable poses among the plethora of possible solutions, for which a scoring function is used.

Another persisting challenge, which is to different extents addressed by protein-protein docking programs, is protein flexibility.(11-14) Molecular dynamics (MD) can account for conformational changes needed for binding at different levels. On a smaller scale, i.e. at the level of atoms, sidechains, loops, small molecules or interfaces, MD is commonly applied to refine docked complexes with the aim of improving their quality.(15-21) MD can be used at a more extensive level, where the docking process is simulated, however modelling spontaneous association and dissociation of proteins is very rare unless coarse-grained models or enhanced sampling methods are used.(22-25) With regard to all-atom MD simulations, enhanced sampling techniques like Markov states models,(26-29) umbrella sampling combined with replica exchange MD, (30-35) elastic-network approaches(35), string method(36), metadynamics(37,38) and other methods have been used to sample conformational change prior to or during binding and to facilitate such binding events under the condition that the binding interface is known(39). Such simulations also present a great opportunity to evaluate binding affinities of known protein-protein complexes. Perthold and Oostenbrink developed GroScore(40) using nonequilibrium free-energy calculations in explicit solvent to score 22000 protein complexes from several Critical Assessment of PRedicted Interactions (CAPRI)(41) sets.(42,43) Kingsley et al.(44) used steered MD and Potentials of Mean Force (PMF) to distinguish between native and non-native poses in 10 docked complexes from ZDOCK.(45) Thirty-nine docked complexes generated by HADDOCK2.2 webserver(2) were ranked by Simões et al.(46) with Molecular Mechanics-Poisson Boltzmann Surface Area (MM-PBSA)(47) calculations. Takemura et al.(48) developed evERdock, that uses the energy representation (ER) method(49) to approximate free-energies of binding and distinguish native from non-native docking models already after only 2ns of MD in explicit solvent.

Recently, alternative and computationally cheap approaches using machine learning (ML) algorithms have been developed to score protein-ligand(50-54) and protein-protein(55-58) complexes, as well as to detect binding interfaces(59-63). Geng et al.(64,65) developed a scoring function (iScore) that combines random walk graph kernel (GraphRank) score with HADDOCK energetic terms. DOcking decoy selection with Voxel-based deep neural nEtwork (DOVE) approach was created by Wang et al., which uses a convolutional deep neural network for evaluating protein docking models and is also available as webserver.(66) Ballester et al. used a Random Forest algorithm with 36 features to predict the binding affinities of protein-ligand complexes.(67,68) Yet another popular method that assesses quality of protein models is Voronoi tessellation-based method(69-71). Olechnovič and Venclovas developed VoroMQA(72), also available as webserver, which uses contact areas derived from Voronoi tessellation of protein structure and tested their approach on Critical Assessment of Structure Prediction (CASP)(73) data set. In supervised machine learning, algorithms first learn from labelled data so that they can apply the learned correlation to new data and predict their labels. There are various classifiers, for example K-nearest Neighbours, Support Vector Machine and its variations, Naïve Bayes or Decision Trees, which can be merged into Random Forest. However, despite of all these efforts, correct and consistent identification of near-native docked models of biomolecular complexes remains a challenge(74,75).

In this work we address the scoring problem by combining standard MD simulations and machine learing to differentiate native from non-native models of protein-protein complexes. We selected 25 complexes from the Docking Benchmark Version 5(76) docked using a local version of HADDOCK2.4 for which the default HADDOCK score was not consistently able to correctly identify on top the models closest to the reference structure. For each complex we selected four models from two top scoring clusters corresponding to near-native and wrong solutions. These, together with the reference crystal structure of the complex, were simulated in explicit solvent for 100 ns each (combined total of 48 ms for all models, references and their replicate simulations). The resulting trajectories were analysed, and eight features, including CAPRI criteria calculated with respect to the starting model, were extracted to build the machine learning model afterwards.

These properties are studied by comparison to both the known reference crystal structure and the starting conformation used MD. Properties from multiple trajectory stretches of 10 and 20 ns were extracted, normalized and fed into a Random Forest (RF) classifier created with the scikit-learn library.(77) The RF classifier was trained on sets of 20 protein-protein complexes and subsequently tested and validated on independent sets of five complexes.

## MATERIALS AND METHODS

### Dataset

230 complexes were chosen from the Docking Benchmark Version 5 (BM5)(76) and docked with HADDOCK version 2.4(1), applying restraints derived from the true interface (ambiguous restraints based on residues making contacts within a 3.9 Å cutoff). These ambiguous restraints have the property to bring the interfaces together without predefining their exact orientation. From these complexes, 25 (13 of which having the top ranked cluster as a non-native model) were selected for our MD approach. Two training and validation sets were defined. Both have as common complexes in the training set the following 15 systems: 1BJ1, 1BUH, 1E4K, 1E6J, 1EAW, 1JPS, 1KAC, 1NSN, 1OC0, 1PXV, 1RV6, 1VFB, 2A5T, 2NZ8 and 3EO1. Training set 1 has in addition five proteins: 1GPW, 1KTZ, 2OOB, 3S9D, and 4G6M. Complexes 1YVB, 2HRK, 2O8V, 2Z0E and 3F1P form the independent test set. In order to assess the impact of the exact composition of the training set on the performance of the RF classifier, in the second set, training set 2, those two sets of five complexes were swapped (from training to test and vice-versa). Complexes with two best scoring clusters were selected where one yielded models of high quality (native) while the other contained mostly incorrect models (non-native). From these clusters the four best models per cluster were selected for MD simulations. In addition, four replicas of the reference (experimental) structure were run for all systems but 1YVB, 2HRK, 2O8V, 2Z0E, and 3F1P, which were added at a later stage. Complexes were selected based on criteria such as complete loops at the protein-protein interface and no ions or co-factors present to avoid any issue with those during MD simulations. Missing side-chains and loops outside the interface were built using MODELLER version 9.12.(78,79) Various quality measures and scores are shown for the native and non-native complexes in Table 1 (see also **Error! Reference source not found**.).

**Table 1:**
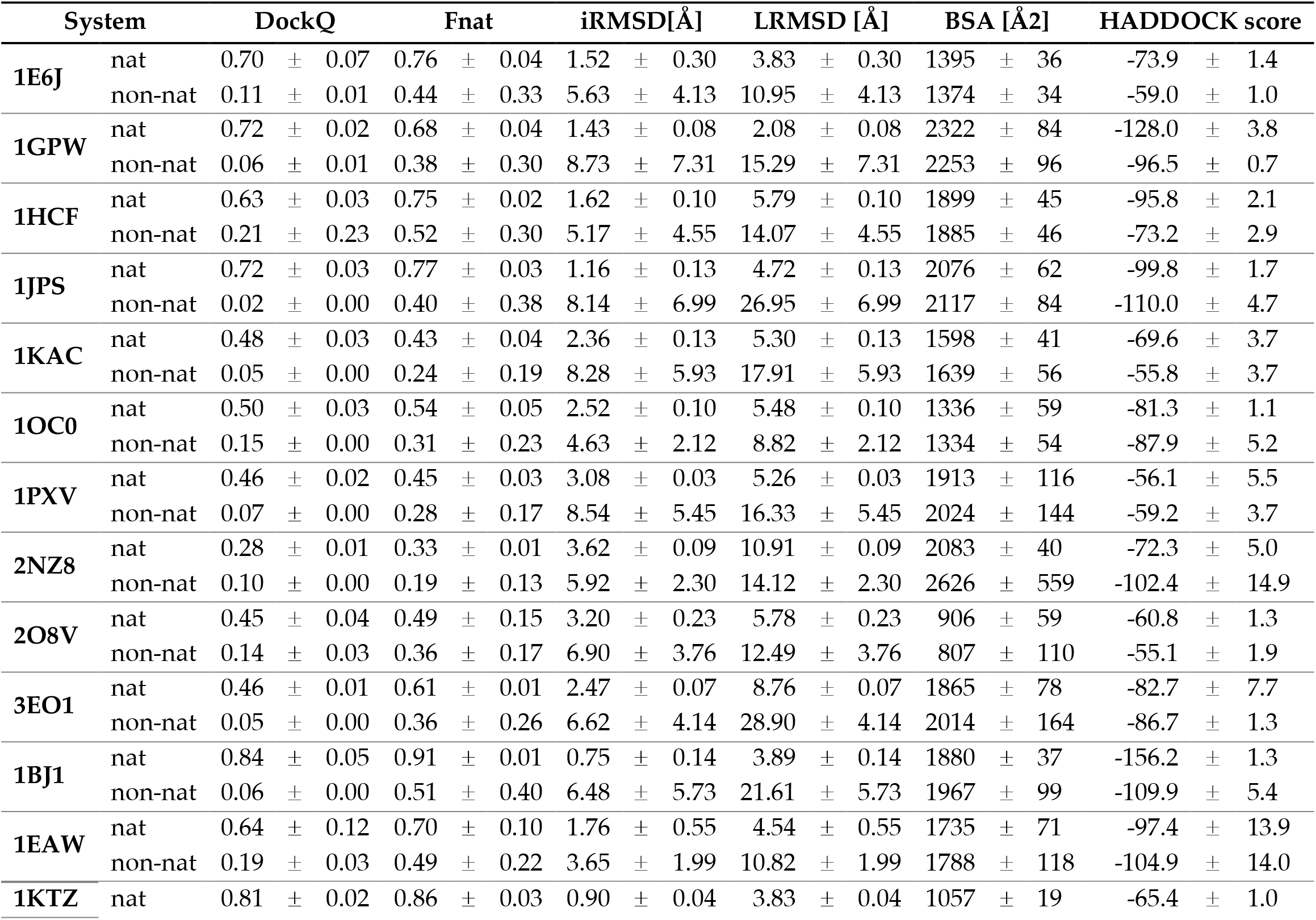
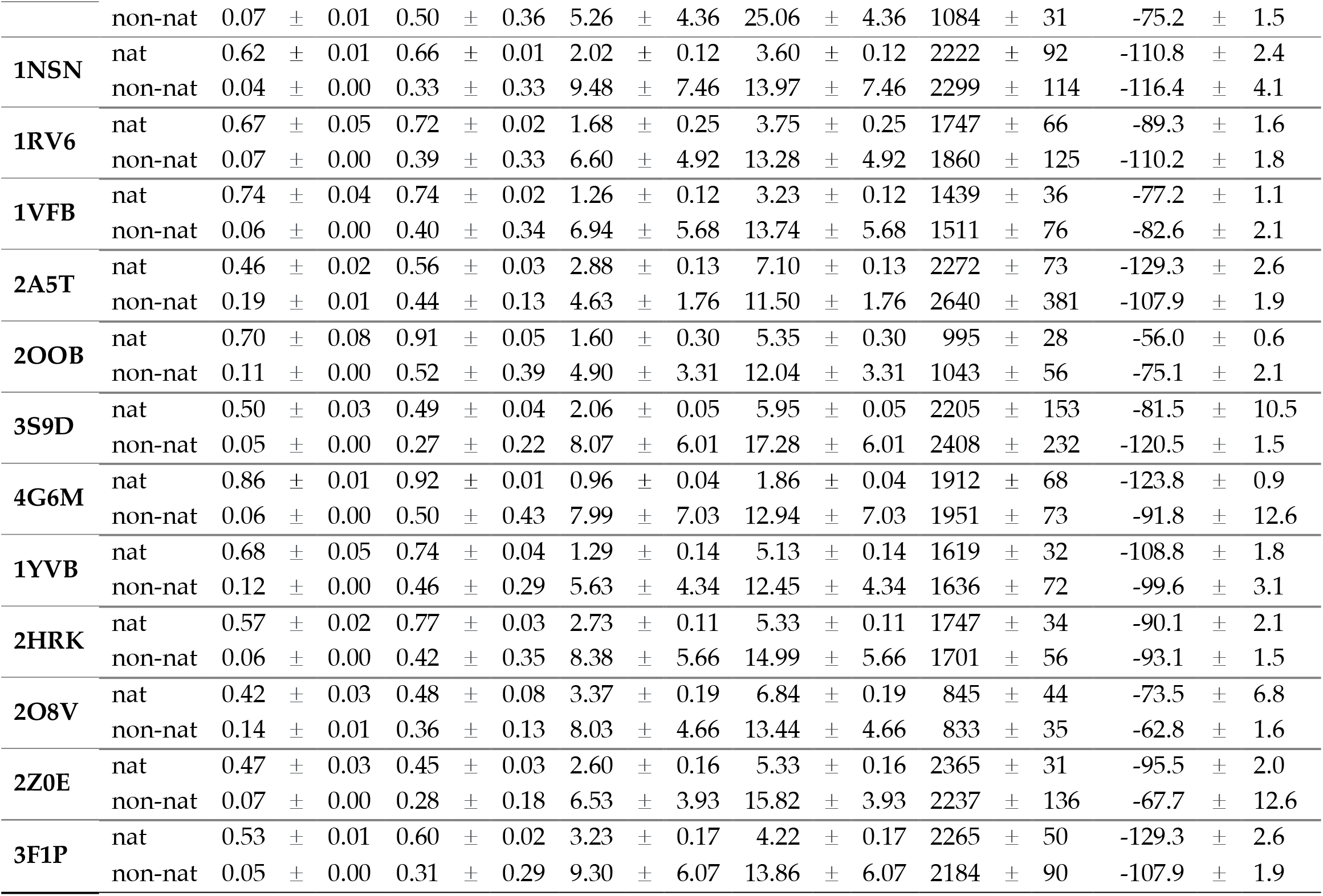
Average properties of top 4 models of the native and non-native clusters selected for further characterisation by MD. The DockQ score (101) was calculated as in the reference(101). Fnat is the fraction of native contacts that takes into account any atom pair forming contacts between proteins within a 5 Å distance cutoff (Fnat=1=100% for the reference). The interface RMSD (i-RMSD) was calculated on the backbone of residues within 10Å of the other protein. The ligand RMSD (l-RMSD) was calculated by fitting on the backbone of the first molecule and calculating the RMSD of the backbone atoms of the second molecule. The buried surface area (BSA) represents the difference of the solvent accessible area of the separated components and the complex. The HADDOCK score is the score used in ranking of HADDOCK models in the final refinement stage (1.0 Evdw(van der Waals intermolecular energy) + 0.2 Eelec(electrostatic intermolecular energy) + 1.0 Edesol(desolvation energy) + 0.1 Eair(distance restraints energy)).

| System |  | DockQ |  |  | Fnat |  |  | iRMSD[Å] |  |  | LRMSD [Å] |  |  | BSA [Å<sup>2</sup>] |  |  | HADDOCK score |  |  |
| --- | --- | --- | --- | --- | --- | --- | --- | --- | --- | --- | --- | --- | --- | --- | --- | --- | --- | --- | --- |
| 1E6J | nat | 0.70 | ± | 0.07 | 0.76 | ± | 0.04 | 1.52 | ± | 0.30 | 3.83 | ± | 0.30 | 1395 | ± | 36 | -73.9 | ± | 1.4 |
|  | non-nat | 0.11 | ± | 0.01 | 0.44 | ± | 0.33 | 5.63 | ± | 4.13 | 10.95 | ± | 4.13 | 1374 | ± | 34 | -59.0 | ± | 1.0 |
| 1GPW | nat | 0.72 | ± | 0.02 | 0.68 | ± | 0.04 | 1.43 | ± | 0.08 | 2.08 | ± | 0.08 | 2322 | ± | 84 | -128.0 | ± | 3.8 |
|  | non-nat | 0.06 | ± | 0.01 | 0.38 | ± | 0.30 | 8.73 | ± | 7.31 | 15.29 | ± | 7.31 | 2253 | ± | 96 | -96.5 | ± | 0.7 |
| 1HCF | nat | 0.63 | ± | 0.03 | 0.75 | ± | 0.02 | 1.62 | ± | 0.10 | 5.79 | ± | 0.10 | 1899 | ± | 45 | -95.8 | ± | 2.1 |
|  | non-nat | 0.21 | ± | 0.23 | 0.52 | ± | 0.30 | 5.17 | ± | 4.55 | 14.07 | ± | 4.55 | 1885 | ± | 46 | -73.2 | ± | 2.9 |
| 1JPS | nat | 0.72 | ± | 0.03 | 0.77 | ± | 0.03 | 1.16 | ± | 0.13 | 4.72 | ± | 0.13 | 2076 | ± | 62 | -99.8 | ± | 1.7 |
|  | non-nat | 0.02 | ± | 0.00 | 0.40 | ± | 0.38 | 8.14 | ± | 6.99 | 26.95 | ± | 6.99 | 2117 | ± | 84 | -110.0 | ± | 4.7 |
| 1KAC | nat | 0.48 | ± | 0.03 | 0.43 | ± | 0.04 | 2.36 | ± | 0.13 | 5.30 | ± | 0.13 | 1598 | ± | 41 | -69.6 | ± | 3.7 |
|  | non-nat | 0.05 | ± | 0.00 | 0.24 | ± | 0.19 | 8.28 | ± | 5.93 | 17.91 | ± | 5.93 | 1639 | ± | 56 | -55.8 | ± | 3.7 |
| 1OC0 | nat | 0.50 | ± | 0.03 | 0.54 | ± | 0.05 | 2.52 | ± | 0.10 | 5.48 | ± | 0.10 | 1336 | ± | 59 | -81.3 | ± | 1.1 |
|  | non-nat | 0.15 | ± | 0.00 | 0.31 | ± | 0.23 | 4.63 | ± | 2.12 | 8.82 | ± | 2.12 | 1334 | ± | 54 | -87.9 | ± | 5.2 |
| 1PXV | nat | 0.46 | ± | 0.02 | 0.45 | ± | 0.03 | 3.08 | ± | 0.03 | 5.26 | ± | 0.03 | 1913 | ± | 116 | -56.1 | ± | 5.5 |
|  | non-nat | 0.07 | ± | 0.00 | 0.28 | ± | 0.17 | 8.54 | ± | 5.45 | 16.33 | ± | 5.45 | 2024 | ± | 144 | -59.2 | ± | 3.7 |
| 2NZ8 | nat | 0.28 | ± | 0.01 | 0.33 | ± | 0.01 | 3.62 | ± | 0.09 | 10.91 | ± | 0.09 | 2083 | ± | 40 | -72.3 | ± | 5.0 |
|  | non-nat | 0.10 | ± | 0.00 | 0.19 | ± | 0.13 | 5.92 | ± | 2.30 | 14.12 | ± | 2.30 | 2626 | ± | 559 | -102.4 | ± | 14.9 |
| 2O8V | nat | 0.45 | ± | 0.04 | 0.49 | ± | 0.15 | 3.20 | ± | 0.23 | 5.78 | ± | 0.23 | 906 | ± | 59 | -60.8 | ± | 1.3 |
|  | non-nat | 0.14 | ± | 0.03 | 0.36 | ± | 0.17 | 6.90 | ± | 3.76 | 12.49 | ± | 3.76 | 807 | ± | 110 | -55.1 | ± | 1.9 |
| 3EO1 | nat | 0.46 | ± | 0.01 | 0.61 | ± | 0.01 | 2.47 | ± | 0.07 | 8.76 | ± | 0.07 | 1865 | ± | 78 | -82.7 | ± | 7.7 |
|  | non-nat | 0.05 | ± | 0.00 | 0.36 | ± | 0.26 | 6.62 | ± | 4.14 | 28.90 | ± | 4.14 | 2014 | ± | 164 | -86.7 | ± | 1.3 |
| 1BJ1 | nat | 0.84 | ± | 0.05 | 0.91 | ± | 0.01 | 0.75 | ± | 0.14 | 3.89 | ± | 0.14 | 1880 | ± | 37 | -156.2 | ± | 1.3 |
|  | non-nat | 0.06 | ± | 0.00 | 0.51 | ± | 0.40 | 6.48 | ± | 5.73 | 21.61 | ± | 5.73 | 1967 | ± | 99 | -109.9 | ± | 5.4 |
| 1EAW | nat | 0.64 | ± | 0.12 | 0.70 | ± | 0.10 | 1.76 | ± | 0.55 | 4.54 | ± | 0.55 | 1735 | ± | 71 | -97.4 | ± | 13.9 |
|  | non-nat | 0.19 | ± | 0.03 | 0.49 | ± | 0.22 | 3.65 | ± | 1.99 | 10.82 | ± | 1.99 | 1788 | ± | 118 | -104.9 | ± | 14.0 |
| 1KTZ | nat | 0.81 | ± | 0.02 | 0.86 | ± | 0.03 | 0.90 | ± | 0.04 | 3.83 | ± | 0.04 | 1057 | ± | 19 | -65.4 | ± | 1.0 |
|  | non-nat | 0.07 | ± | 0.01 | 0.50 | ± | 0.36 | 5.26 | ± | 4.36 | 25.06 | ± | 4.36 | 1084 | ± | 31 | -75.2 | ± | 1.5 |
| **1NSN** | nat | 0.62 | ± | 0.01 | 0.66 | ± | 0.01 | 2.02 | ± | 0.12 | 3.60 | ± | 0.12 | 2222 | ± | 92 | -110.8 | ± | 2.4 |
|  | non-nat | 0.04 | ± | 0.00 | 0.33 | ± | 0.33 | 9.48 | ± | 7.46 | 13.97 | ± | 7.46 | 2299 | ± | 114 | -116.4 | ± | 4.1 |
| **1RV6** | nat | 0.67 | ± | 0.05 | 0.72 | ± | 0.02 | 1.68 | ± | 0.25 | 3.75 | ± | 0.25 | 1747 | ± | 66 | -89.3 | ± | 1.6 |
|  | non-nat | 0.07 | ± | 0.00 | 0.39 | ± | 0.33 | 6.60 | ± | 4.92 | 13.28 | ± | 4.92 | 1860 | ± | 125 | -110.2 | ± | 1.8 |
| **1VFB** | nat | 0.74 | ± | 0.04 | 0.74 | ± | 0.02 | 1.26 | ± | 0.12 | 3.23 | ± | 0.12 | 1439 | ± | 36 | -77.2 | ± | 1.1 |
|  | non-nat | 0.06 | ± | 0.00 | 0.40 | ± | 0.34 | 6.94 | ± | 5.68 | 13.74 | ± | 5.68 | 1511 | ± | 76 | -82.6 | ± | 2.1 |
| **2A5T** | nat | 0.46 | ± | 0.02 | 0.56 | ± | 0.03 | 2.88 | ± | 0.13 | 7.10 | ± | 0.13 | 2272 | ± | 73 | -129.3 | ± | 2.6 |
|  | non-nat | 0.19 | ± | 0.01 | 0.44 | ± | 0.13 | 4.63 | ± | 1.76 | 11.50 | ± | 1.76 | 2640 | ± | 381 | -107.9 | ± | 1.9 |
| **2OOB** | nat | 0.70 | ± | 0.08 | 0.91 | ± | 0.05 | 1.60 | ± | 0.30 | 5.35 | ± | 0.30 | 995 | ± | 28 | -56.0 | ± | 0.6 |
|  | non-nat | 0.11 | ± | 0.00 | 0.52 | ± | 0.39 | 4.90 | ± | 3.31 | 12.04 | ± | 3.31 | 1043 | ± | 56 | -75.1 | ± | 2.1 |
| **3S9D** | nat | 0.50 | ± | 0.03 | 0.49 | ± | 0.04 | 2.06 | ± | 0.05 | 5.95 | ± | 0.05 | 2205 | ± | 153 | -81.5 | ± | 10.5 |
|  | non-nat | 0.05 | ± | 0.00 | 0.27 | ± | 0.22 | 8.07 | ± | 6.01 | 17.28 | ± | 6.01 | 2408 | ± | 232 | -120.5 | ± | 1.5 |
| **4G6M** | nat | 0.86 | ± | 0.01 | 0.92 | ± | 0.01 | 0.96 | ± | 0.04 | 1.86 | ± | 0.04 | 1912 | ± | 68 | -123.8 | ± | 0.9 |
|  | non-nat | 0.06 | ± | 0.00 | 0.50 | ± | 0.43 | 7.99 | ± | 7.03 | 12.94 | ± | 7.03 | 1951 | ± | 73 | -91.8 | ± | 12.6 |
| **1YVB** | nat | 0.68 | ± | 0.05 | 0.74 | ± | 0.04 | 1.29 | ± | 0.14 | 5.13 | ± | 0.14 | 1619 | ± | 32 | -108.8 | ± | 1.8 |
|  | non-nat | 0.12 | ± | 0.00 | 0.46 | ± | 0.29 | 5.63 | ± | 4.34 | 12.45 | ± | 4.34 | 1636 | ± | 72 | -99.6 | ± | 3.1 |
| **2HRK** | nat | 0.57 | ± | 0.02 | 0.77 | ± | 0.03 | 2.73 | ± | 0.11 | 5.33 | ± | 0.11 | 1747 | ± | 34 | -90.1 | ± | 2.1 |
|  | non-nat | 0.06 | ± | 0.00 | 0.42 | ± | 0.35 | 8.38 | ± | 5.66 | 14.99 | ± | 5.66 | 1701 | ± | 56 | -93.1 | ± | 1.5 |
| **2O8V** | nat | 0.42 | ± | 0.03 | 0.48 | ± | 0.08 | 3.37 | ± | 0.19 | 6.84 | ± | 0.19 | 845 | ± | 44 | -73.5 | ± | 6.8 |
|  | non-nat | 0.14 | ± | 0.01 | 0.36 | ± | 0.13 | 8.03 | ± | 4.66 | 13.44 | ± | 4.66 | 833 | ± | 35 | -62.8 | ± | 1.6 |
| **2Z0E** | nat | 0.47 | ± | 0.03 | 0.45 | ± | 0.03 | 2.60 | ± | 0.16 | 5.33 | ± | 0.16 | 2365 | ± | 31 | -95.5 | ± | 2.0 |
|  | non-nat | 0.07 | ± | 0.00 | 0.28 | ± | 0.18 | 6.53 | ± | 3.93 | 15.82 | ± | 3.93 | 2237 | ± | 136 | -67.7 | ± | 12.6 |
| **3F1P** | nat | 0.53 | ± | 0.01 | 0.60 | ± | 0.02 | 3.23 | ± | 0.17 | 4.22 | ± | 0.17 | 2265 | ± | 50 | -129.3 | ± | 2.6 |
|  | non-nat | 0.05 | ± | 0.00 | 0.31 | ± | 0.29 | 9.30 | ± | 6.07 | 13.86 | ± | 6.07 | 2184 | ± | 90 | -107.9 | ± | 1.9 |

### Molecular Dynamics simulations

The selected complexes were simulated and subsequently analysed using the GROMACS simulation package(80) version 2019 and the CHARMM36m(81) forcefield. Protein structures were first optimized in vacuum using the steepest-descent algorithm (up to 5000 steps) and subsequently solvated in a rhombic dodecahedral box of TIP3P(82) water. Minimal solute to box distances were set to 1.4 nm, and sodium and chloride ions were added to neutralise the box and reach a concentration of 150 mM. A second optimization stage was performed (up to 25000 steps if convergence – standard settings – was not reached before). A first MD run of 50 ps was then performed at 50 K in the NVT ensemble using a velocity-rescaling thermostat(83,84) with a 0.1 ps time constant. Initial velocities were randomly assigned according to a Maxwell–Boltzmann distribution at 50 K and the system was subsequently heated up to 150 and 300 K. During this equilibration phase, all heavy atoms of the proteins were positionally restrained using a decreasing force constant of 1000 (50 K), 100 (150 K), and 10 (300 K) kJ mol^−1^ nm^−2^ in x-, y-, and z-coordinates. Production runs of 100 ns in length were performed in an NPT ensemble using the Berendsen barostat(84) with isotropic pressure scaling, a time constant of 1 ps and an isothermal compressibility of 4.5 × 10^−5^ bar^-1^. For all MD simulations the leapfrog integration scheme(85) with a timestep of 2 fs was used and covalent bonds were restrained using the LINCS algorithm(86). Neighbour searching was performed using a Verlet-based cut-off scheme updated every 10 steps with a cut-off of 1 nm. For the Van der Waals interactions a twin range cut-off with a smooth switch to zero between 1 and 1.2 nm was used. The Particle Mesh Ewald method(87) was used to calculate the long-range electrostatics. The total simulation time for all complexes sums up to 48 ms. Trajectory frames were written to disk every 500 ps for further analysis.

### Analysis

The definition of fraction of common contacts, interface residues and ligands followed the CAPRI classification.(88) Intermolecular contacts were considered as any atom pair between proteins within 5 Å, and the contacts evolution as a function of simulation time was calculated using the *gmx hbond -contact* analysis tool. Interface residues are defined as residues, with at least one atom within 10 Å of the other molecule, and interface RMSD (i-RMSD) was calculated on the backbone of these residues. Ligand RMSD was calculated by fitting on the backbone of the first molecule and calculating the RMSD of the backbone atoms of the second molecule. The DockQ values reported in Table 1 are based on a combination of all of these properties (Fnat, i-RMSD, l-RMSD) as described in reference.(89) Distance between centres of mass of proteins, hydrogen bonds, buried surface area were calculated using the standard GROMACS analysis tools (*gmx distance, gmx hbond* and *gmx sasa*). The interaction energy terms were calculated as the sum of short-range Coulombic and Lennard-Jones interactions between the two proteins or between the proteins and water. To compensate for the varying size of the systems and interfaces for further machine learning analysis, all properties were evaluated relatively to their values at the beginning of the trajectory. Hence, only changes of the properties over time are observed, not their absolute numbers. These time series were extracted for all trajectories and were used to feed the Random Forest classifier.

### Random Forest model for native vs non-native classification

To select the most accurate classifier for our task, we evaluated the performance of the following set of binary classifiers available in the Scikit-learn library(77): gnb = Gaussian navie bayes, KNN = K Neighbors Classifier (n_neighbors=1), MNB = Multinomial naïve bayes, BNB = Bernoulli naïve bayes, LR = Logistic Regression, SDG = stochastic gradient descent Classifier, SVC = Support Vector classification, LSVC = Linear SVC, NSVC = Nu SVC, RF = Random Forest Classifier (see Figure S7). These classifiers were tested on the training set every 10 ns, using repeated K-fold cross-validation with 10 splits and 10 repeats with a test size of 25% (i.e. 5 complexes). Since Random Forest showed the highest accuracy throughout the entire trajectory, it was selected for our approach. Random Forest combines multiple components of randomness: First, the training set is divided into multiple bootstrapped copies and predictions from these are aggregated, which reduces the variance, or overfitting compared to individual decision trees. Moreover, to further decrease the correlation among trees, a random subset of features is selected at each tree split. In this work the Scikit-learn library(77) 0.23.1 was used again to create the Random Forest classifier. The Grid Search algorithm was used to search for the optimal parameters based on the last 20 ns of trajectories of training set 1, as described in the Dataset part of the Methods section. The following parameters were used to create the Random Forest classifier: Number of trees in the forest set to 1000, bootstrap samples used when building trees, square root of features selected at each split, maximum depth of the tree set to 50, minimum number of samples required to split an internal node set to 2, minimum number of samples required to be at a leaf node set to 1. The model was subsequently trained on different 20 ns patches of the trajectory. Average receiver operating characteristic (ROC) curve, accuracy and f1-score were calculated from 10 times 10-fold cross-validation optimizations. Accuracy is calculated as the fraction of correct predictions (true positive (TP) + true negative (TN)) out of the total number of predictions (TP + TN + false positive (FP) + false negative (FN)). F1-score is calculated as: TP / (TP + ½ (FP + FN)). True positive rate (TPR) also known as sensitivity are the fraction of TP out of the positives and true negative rate (TNR) and the fraction of TN out of the negatives. Relation between these two metrics is reflected in the ROC curve.

## RESULTS

Twenty-five protein-protein complexes from BM5(76) modelled with HADDOCK using true interface information (but no specific contacts) were selected on the basis that their two best scoring clusters showed models of utterly different quality (near native and non-native). For about half of those complexes (13 out of 25) HADDOCK ranks on top a non-native model. Our aim was to use MD simulations complemented by machine learning to distinguish native from non-native models. The quality of all models, together with the buried surface area and HADDOCK score are listen in **Error! Reference source not found**. (see also **Error! Reference source not found**.).

### Standard MD

For each complex, the four best scoring models per native and non-native clusters were selected and simulated, each with two replicas per model, for 100 ns using GROMACS. Additionally, for the 20 complexes of training set 1, simulations of the reference structure (crystal structure; four MD replicas per reference structure) were performed and analysed. Properties of all complexes and models were measured with respect to both the reference crystal structure of the complex and the docking model used as starting conformation for MD simulations. For training the machine learning model, relative values with respect to the start of the production run (ref-orig) were used as input, which partly normalizes the differences due to the varying size of the complexes.

### Can MD improve the quality of the models?

The quality of protein-protein complexes is commonly assessed by comparing a number of properties to their reference (usually, experimental) structure. In CAPRI, those properties are ligand-RMSD, interface-RMSD and fraction of native contacts.

In the first part of our work, we looked at these properties during the course of the MD trajectories of both the HADDOCK models and the reference crystal structures and compared them to the experimental crystal structure of the complex. Figure 1 shows the distributions of l-RMSD, i-RMSD and Fnat with respect to the respective reference crystal structures for all the 20 complexes of training set 1 and their evolution over different stretches along the simulation. This analysis is based on a total of 160 simulations of both native and non-native clusters, and 80 simulations of the reference crystal structures, each of 100 ns in length.

**Figure 1:**
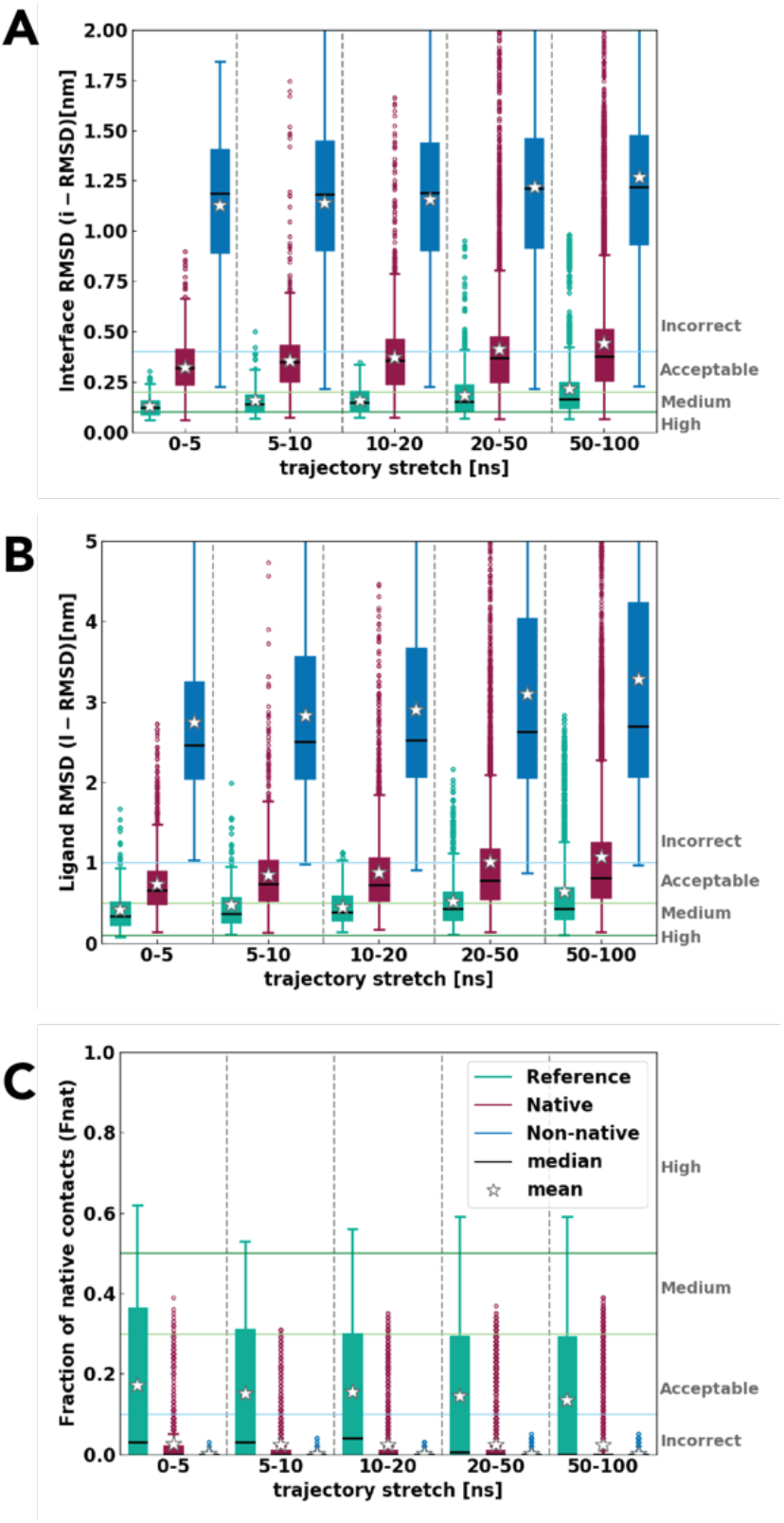
A) Interface RMSD B) Ligand RMSD and C) fraction of native contacts for native, non-native and reference structures from the complex crystal structure for all 20 complexes. The boxplot shows the interquartile range with its median as black lines, mean as stars, whiskers in error bars and outliers in circles. Reference in green, native clusters in burgundy, non-native in blue.

As expected, all systems including those started from the reference crystal structures undergo changes along the course of the simulation. The magnitude of these changes is, however, somewhat surprising: Already in the first 5 ns of the production runs, even the reference structures lose on average up to 80% of their original contacts (Figure 1 C). This might be due to the initial rearrangement of the residues during the heating up phases of the simulation and relatively tight definition of intermolecular contacts (5 Å). Indeed, with regard to the simulation of the reference structures, we noticed in all cases only small changes in the conformation of the backbone at partners interface (Figure S2). All three groups of complexes deviate further from the crystal structure over time, which results in poorer model quality towards the end of the simulation. In particular, while the reference simulations remain during the entire trajectory within the acceptable quality CAPRI category, near-native docked models reach the incorrect area by the end of the simulation, while non-native models never reach the correct category. Despite this, a clear and consistent separation of native and non-native models is evident throughout the entire course of the simulation for all three properties reported in Figure 1. While 100 ns of MD simulation is not able to improve the quality of the initial models, it clearly allows to differentiate between near-native and non-native models based on a comparison to the known crystal structure.

### Can MD distinguish between native and non-native complexes based on CAPRI measures?

The second question we wanted to address in our work is if standard MD is able to capture any different behaviour of native and non-native models. This is particularly important, because in a realistic scenario there will be no reference structure available. To assess this, we selected for each simulation the model at the starting point of the production run as a reference (hereafter ref-orig). Figure 2 depicts the same properties as Figure 1 which we denote by an additional “-orig” label to distinguish from the same values with respect to the reference crystal structure (l-RMSD^orig^, i-RMSD^orig^ and Fnat^orig^), so as to highlight their behaviour relatively to the initial binding mode (ref-orig). For both ligand and interface RMSD^orig^, the reference crystal structures show the lowest values, pointing to the (expected) higher stability of the experimental complexes compared to their docked models. More interestingly, the near-native complexes show overall higher stability (less deviations from the initial values) than the non-native ones. Even though the distributions of both RMSD^orig^s are largely overlapping, their means are clearly distinguishable at the end of the simulation (∼0.5 nm for i-RMSD^orig^ and ∼1 nm for l-RMSD^orig^ difference from ref-orig (Figure 2 A and B)). Importantly, these differences increase over the simulation time most often due to the increasing instability of non-native complex configurations in the second half of the simulations (Figures S2, S3). A similar trend is seen in the decrease of Fnat^orig^ over time (Figure 2 C, Figure S4). While the reference and native structures lose up to 70% and 80% of initial contacts, respectively, the non-native models lose almost all of them towards the end of the simulation.

**Figure 2:**
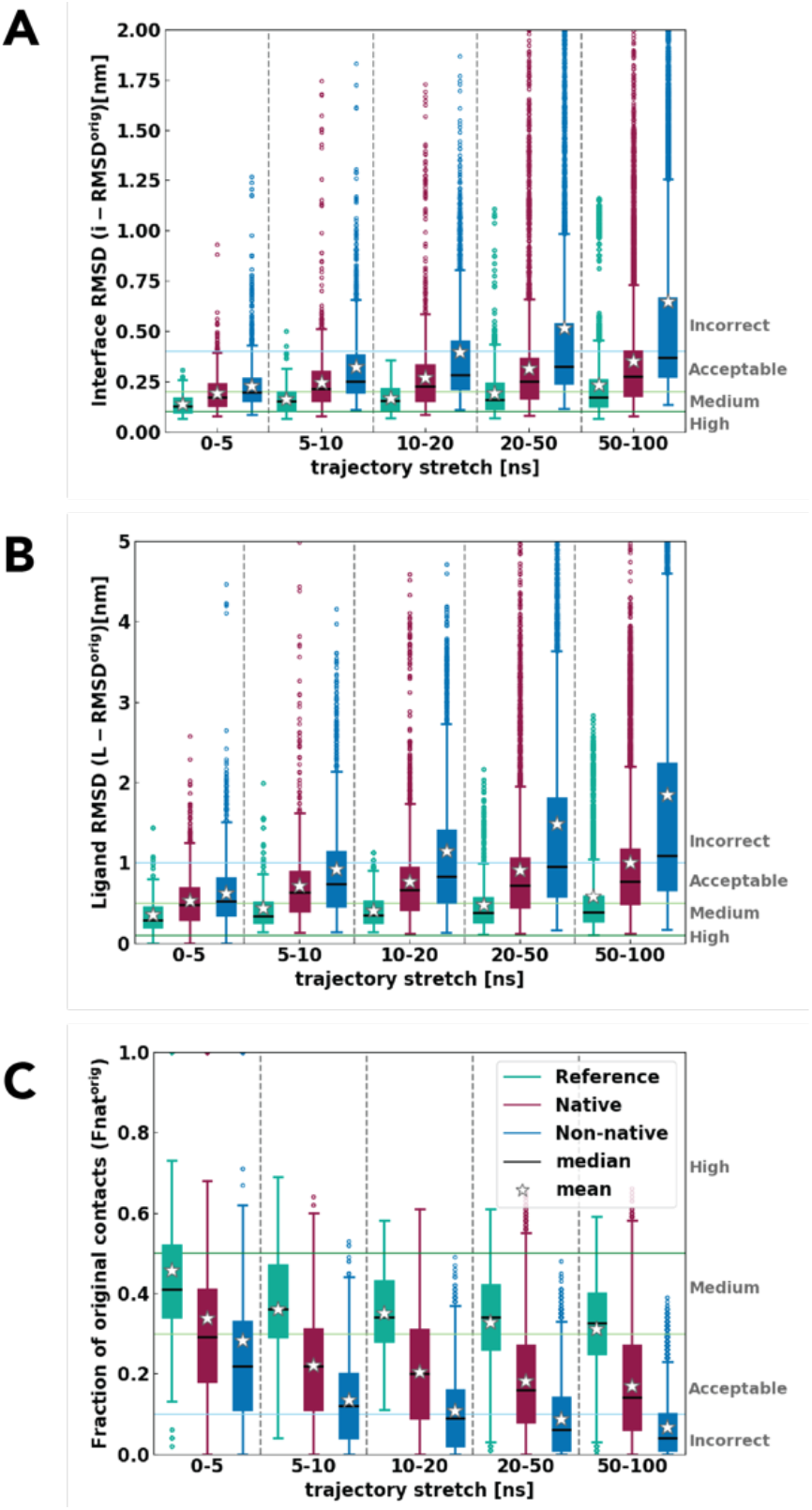
A) Interface RMSD^orig^ B) Ligand RMSD^orig^ and C) fraction of native contacts (Fnat^orig^) for native, non-native and reference crystal structures with respect to the beginning of the trajectory for all 20 complexes. The boxplot shows the interquartile range with its median as black lines, mean as stars, whiskers and outliers in circles. Reference in teal, native clusters in burgundy, non-native in blue.

Interestingly, the trends are evident already within the first 5 ns of the simulation, where even the reference simulations lose up to 50% of original contacts. This implies than even such a short stretch of plain MD causes significant changes in the protein interfaces, as already highlighted by Figure 1. For a comparison, CAPRI criteria are also indicated in Figure 2. These should help to illustrate the order of structural changes occurring at the interface and the rearrangement of the complex during MD. In all three properties non-native models enter the incorrect category in the second half of the simulation (e.g., after 50 ns), while reference and native models remain in the acceptable to medium quality area (**Error! Reference source not found**. -**Error! Reference source not found**.).

### Comparison of other structural and energetic properties

A number of additional properties were measured for selected complexes. **Error! Reference source not found**. A and B show the evolution of the buried surface area and of the distance between the COMs of the two binding partners. While for reference structures both properties are relatively stable throughout the simulation, for non-native models the BSA is consistently decreasing while the distance between the centre of masses increases. By the end of the trajectory, the variations amount on average to 5 nm^2^ and 0.3 nm, respectively. The number of hydrogen bonds is as well slightly decreasing (**Error! Reference source not found**. C) for the non-native complexes. An identical behaviour can be seen in the non-bonded interaction energies between proteins and proteins and water depicted in **Error! Reference source not found**. D and E. As expected, due to the initial (generally unfavourable) conformation of non-native complexes, the interaction between protein partners becomes weaker with increasing simulation time for these models. This is compensated by more favourable interactions with water, which can be seen in the decrease of the non-bonded interaction energy. The native complexes don’t deviate considerably from their reference complexes, which is consistent with previous findings and CAPRI properties. Such a clear distinction in the behaviour of native and non-native complexes during standard MD of 100 ns or less is rather remarkable. We therefore decided to exploit the measured properties to develop a machine learning model that could help us classify models as native or non-native based on their simulation properties.

### Random Forest classifier

#### Cross-validation on training set 1

Properties calculated from simulations of 20 HADDOCK models of training set 1 were mixed and labelled as native and non-native, based on the quality of the initial model they were extracted from. They were divided into trajectory stretches of 20 ns. An example of such distribution of all properties and their scatter plots in the last 20 ns of the trajectory is shown in **Error! Reference source not found**.. To choose the most fitting classifier for our purposes, the trajectory was divided into stretches of 10 ns and, for each of this time frames, a number of classifiers from the Scikit library were trained. Ten ns were used to obtain a more detailed overview of ML classifying accuracy along the trajectory. The accuracy score for all of them are summarised in **Error! Reference source not found**.. The Random Forest classifier reached clearly higher accuracy than the other ones and was chosen for further classification of the complexes. Random Forest combines bootstrapping of the training set with random feature selection at each tree split. First, the search for optimal parameters was performed using the Grid Search algorithm based on the last 20 ns of trajectories from the training set. The best parameters were selected, which are listed in the Methods section. Subsequently the model was trained considering different trajectory stretches. Its accuracy was calculated within its own cross-validated training set 1. On top of the inherent randomness that Random Forest involves, stratified K-fold cross-validation with 100 splits was used. Accuracy, precision, recall, f1-score, area under curve (AUC) of Receiver Operating Characteristic (ROC), calculated for 5 trajectory stretches of 20 ns are summarised in Table 2. The accuracy of the RF classifier starts at 0.77 and reaches 0.86 in the last 20 ns of the simulation. Similar trends can be seen in all performance metrics. Interestingly, after 40-60 ns all metrics start converging. RF classifiers allow to evaluate the importance of the various features used in training. **Error! Reference source not found**. summarises features importance assessed on the last 20 ns of the trajectory of training set 1. Changes in i-RMSD^orig^, BSA^orig^ and Fnat^orig^ are the most important in distinguishing native from non-native models, while the number of hydrogen bonds shows the smallest importance. However, overall, the relative differences between property importance were not very substantial.

**Table 2:**
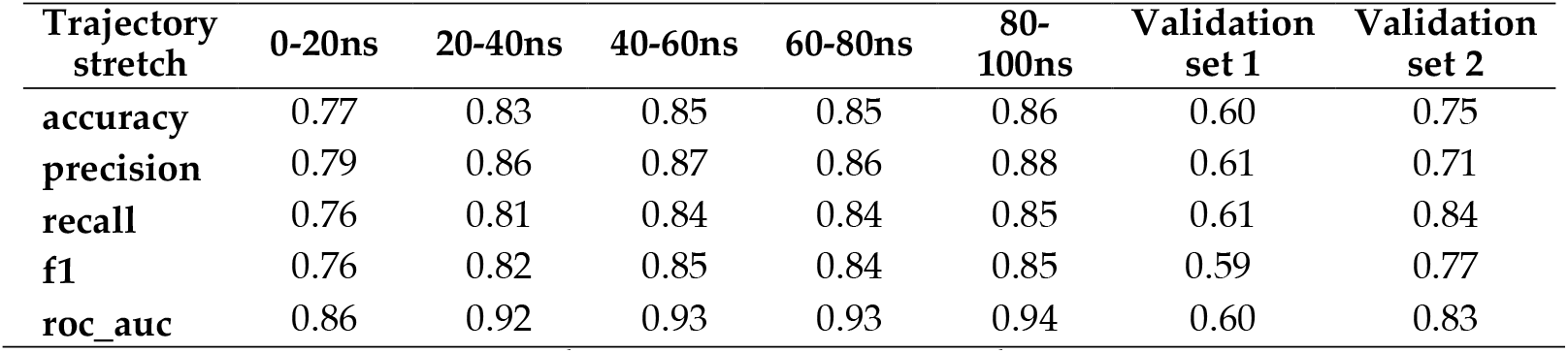
Scoring performance of the RF classifier based on the cross-validated training set 1 and both validation sets.

#### Validation on independent validation sets

While the RF classifier performed well on the cross-validated training test alone, a fair comparison would be to test its accuracy on an external validation test. To this end, five additional complexes were selected (validation set 1), and their native and non-native models were simulated for 100 ns as for the training set 1 (see Methods). CAPRI properties of the validation set throughout the trajectory are shown in **Error! Reference source not found**.. Surprisingly, here the differences between native and non-native complexes were not as high as in the training set. When the model was tested on the independent validation set, its accuracy and f1-score reached 0.60 and 0.59 respectively (Table 2). This is significantly lower than for the original training set. The same is observed for the AUC with 0.60 against 0.93 for the training set (Figure 3B). Prompted by this finding, we performed another round of training and validation of the model using a different distribution of complexes between training and test sets (training set 2 and validation set 2) (see Methods). For this, five randomly selected complexes from the original training set were swapped with the validation set. After training on this second data set, the RF classifier shows a better performance at distinguishing native from non-native models for the validation set with accuracy and AUC of 0.75 and 0.83, respectively (Table 2 and Figure 3C). This implies that the model accuracy depends on the nature of the initial complexes and their stability during MD. But even in the unfavourable scenario where the behaviour of both classes of complexes was rather similar (validation set 1), our model was able to correctly identify native complexes with accuracy of 60%.

**Figure 3:**
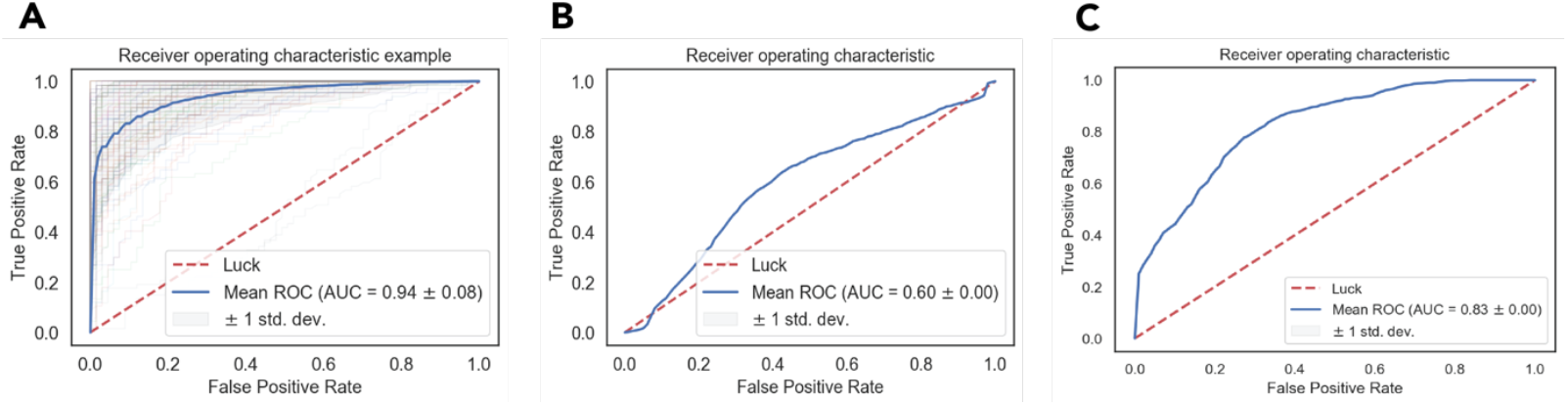
Receiver operating characteristic (ROC) curves showing: A) training set 1 using stratified K-fold cross-validation split 100 times. Individual splits shown in thin lines and their mean in blue bold line. B) Validation set 1 C) Validation set 2

## DISCUSSION

The flexible nature of proteins and their interfaces naturally evolve into dynamic interactions among binding partners.(90-92) Our work makes use of such dynamics to tackle the intricate task of correctly scoring models of protein-protein complexes obtained by docking. There are not many MD approaches known to focus on this task without using enhanced sampling or free-energy calculations. In this work native and non-native models of 25 complexes from the docking benchmark 5 docked with HADDOCK2.4 and their reference crystal structures were simulated in multiple copies, each for 100 ns in length (cumulative time: 48 μs). The trajectories were analysed, and their properties used to feed a Random Forest classifier.

Running MD simulation on docked models could lead to a spontaneous complex rearrangement to a more favourable position (aka induced fit) provided sufficient sampling. Here, by comparing simulated HADDOCK models and their structures at the beginning of the production run (ref-orig) to the original reference crystal structure we first assessed if MD simulations would allow to improve the quality of the models. Significant changes were observed at the interface of all simulated structures, even the crystal structure. However, little to no improvement was observed for the near-native models. Perhaps, much longer time scales would be necessary to capture spontaneous rearrangements of protein complexes, similar to long plain MD simulations which have been used to observe domain rearrangement in single proteins.(93-95) Pan et al.(22) observed reversible binding and unbinding of multiple protein complexes in time scale of hundreds of microseconds using the tempered binding approach and dedicated hardware built in house. In that work the native binding was not reached by sampling of the entire protein surface while proteins stayed in close contact, but rather after a repeated dissociation and reassociation of the complex (mimicked by docking here). This observation was analogous to ours, since we also did not observe any rearrangement from non-native to native binding pose while the proteins stayed in contact. One can assume that protein-protein reassociation would be needed in our case too, however this can hardly be achieved in the 100 ns timescale of our simulations. Nonetheless, the simulations revealed a clear difference in the behaviour of non-native and native models or reference during that timescale.

In the second part of our work, we examined in a realistic scenario, i.e. in the absence of a reference structure, if MD simulations could be used as a scoring tool to distinguish native from non-native models. For this, properties of the simulations were compared to their values at the beginning of the trajectory, measuring thus changes with respect to the original starting model. Here, clearly the crystal structures followed by the native models exhibited the highest stability in contrast to the non-native models. This is a promising observation, where, without any prior knowledge of model quality, one can see differences in their stability during simulation which allow to point out the less stable / non-native models. Radom et al.(96) could similarly distinguish decoys from native structures based on their stability during MD and noticed a few exceptions, where the wrong binding pose would find the correct conformation throughout the simulation. Akin cases were seen in our study as well, nonetheless they were statistically not significant enough to influence the overall trend. Prevost et al.(39) used a similar MD ranking approach for docked complexes. They suggested to alter the CAPRI scoring criteria and take the dynamic properties (l-RMSD, i-RMSD and Fnat) into account by comparing to simulations of the reference crystal structures of the complexes. To correlate system properties to the simulated reference, instead of its crystal structure, could be advantageous, yet in our case somewhat redundant since were already able to observe stability differences even without the reference present.

Based on these findings a machine learning model was developed to classify complexes as native or non-native in an automated manner rather than by visual inspection of the properties as a function of simulation time. The performance of the model was assessed on cross-validated training sets as well as two independent validation sets. The accuracy on the training set reached 0.85 and ranged between 0.60 to 0.75 for validation sets. While the accuracy on the training set was increasing as a function of the simulation time window considered, it seems to converge after 50 ns. This would indicate that shorter simulations of 50 ns might already be sufficient for this kind of classification in the future.

Such combination of docking, molecular dynamics simulation and machine learning as used in this work, are becoming more common.(97,98) Still their application to protein-protein interactions remains limited. A number of studies have focused on scoring docked protein-protein models neglecting their dynamic nature. Pfeiffenberger et al.(56) applied an Extremely Randomized Tree Classifier to rank poses generated by SwarmDock(99,100) for 11 CAPRI targets classified by their ligand RMSD. Using 109 molecular descriptors, they obtained accuracy between 0.6-0.7 at distinguishing native from non-native complexes with residue– residue contact descriptors, which is comparable to our results. Das and Chakrabati(101) employed a Support Vector Machine to differentiate between native and non-native interfaces using features like accessible, buried surface area and frequency of salt bridges or hydrogen bonds. F1-score of 0.8 was achieved on an external dataset at distinguishing between native and non-native models. Both studies found that the key features for protein-protein interactions are intermolecular contacts and accessible/buried surface area. Comparable accuracies (0.7-0.9) were obtained by the DOVE approach (66) on validation and training sets. iScore(64) as a graph-kernel based scoring method ranked amongst the top scoring approaches on the CAPRI scoring set. Our MD-based scoring method not only reaches a state of art techniques level of accuracy but also introduces a novel way to incorporate information about the dynamics of protein complexes into scoring. The encouraging results obtained show that, even if the behaviour of the complexes remain similar for both native and non-native models, the RF classifier is able to distinguish them in up to 75% of the cases. From the feature importance analysis, the most important properties for classifying the quality of a model are suggested to be the fraction of native contacts, interface RMSD and buried surface area with respect to the starting model.

## DATA AVAILABILITY

All MD trajectories without water and their topologies are deposited in zenodo (http://doi.org/10.5281/zenodo.4629895). GROMACS scripts, extracted features and jupyter notebooks with Random Forest classifiers are available in the Github repository (https://github.com/haddocking/MD-scoring).

## SUPPLEMENTARY DATA

Supplementary Data are available at NAR online.

## ACKNOWLEDGEMENT

We thank the entire computational structural biology group at Utrecht University for fruitful discussions, and in particular Dr. Francesco Ambrosetti for his input on machine learning. This research used the Savio computational cluster resource provided by the Berkeley Research Computing program at the University of California, Berkeley (supported by the UC Berkeley Chancellor, Vice Chancellor for Research, and Chief Information Officer).

